# Trendy: Segmented regression analysis of expression dynamics for high-throughput ordered profiling experiments

**DOI:** 10.1101/185413

**Authors:** Rhonda Bacher, Ning Leng, Li-Fang Chu, James Thomson, Christina Kendziorski, Ron Stewart

## Abstract

High throughput expression profiling experiments with ordered conditions (e.g. time-course or spatial-course) are becoming more common for profiling detailed differentiation processes or spatial patterns. Identifying dynamic changes at both the individual gene and whole transcriptome level can provide important insights about genes, pathways, and critical time-points. We present an R package, **Trendy**, which utilizes segmented regression models to simultaneously characterize each gene’s expression pattern and summarize overall dynamic activity in ordered condition experiments. For each gene, Trendy finds the optimal segmented regression model and provides the location and direction of dynamic changes in expression. We demonstrate the utility of Trendy to provide biologically relevant results on both microarray and RNA-seq datasets. Trendy is a flexible R package which characterizes gene-specific expression patterns and summarizes changes of global dynamics over ordered conditions. Trendy is freely available as an R package with a full vignette at https://github.com/rhondabacher/Trendy.

## Background

High throughput expression profiling technologies have become essential tools for advancing insights into biological systems and processes. By profiling over ordered conditions such as time or space, the power of microarrays and sequencing can be further leveraged to study the dynamics of biological processes. Of great interest in time-course or spatial-course experiments is to identify genes with dynamic expression patterns, which can provide insight on regulatory genes (1) and highlight key transitional periods (2).

Many methods for time-course experiments are aimed at identifying differentially expressed genes across multi-series time-courses and are detailed in a review by Spies and Ciaudo, 2015 (3). Here we focus on analyzing single-series time-course experiments, where one can identify dynamic genes via each genes’s expression path or pattern across time.

Single-series time course methods have largely focused on the clustering of gene expression (4; 5), which can be used to construct regulatory networks, and they do not typically emphasize the information provided by each gene’s individual expression path. EBSeqHMM (6) was developed in part to address this deficiency and employs a hidden Markov model to classify genes into distinct expression paths. Despite its utility, differences between time-points may not be sufficiently detectable for extensive or densely sampled time-course experiments with subtle expression changes. FunPat (7) is another method which can analyze a single-series time course and does so based on comparing changes in expression to a user-specified baseline. It assumes genes with a common biological annotation have a common expression pattern and is able to link gene sets to expression patterns. Although the patterns are represented visually and searchable based on biological annotation and pattern ID, they are not easily searched by the user based on characteristics of their trend as they are not summarized in this way. EBSeqHMM requires at least two replicates per time-point, while FunPat requires only one time-point to have at least two replicates. However, in long or dense time-courses replicates may be missing if they are sacrificed for an increased sampling rate.

We have developed an R package, Trendy, which employs segmented regression models to simultaneously characterize each gene’s expression pattern and summarize overall dynamic activity in time-course experiments. For each gene, Trendy fits a set of segmented regression models with varying numbers of breakpoints. Each breakpoint represents a dynamic change in the gene’s expression profile over time. A model selection step then identifies the model with the optimal number of breakpoints. Trendy implements functions to visualize and order the top dynamic genes and their trends. The top dynamic genes are defined as those that are well-profiled based on the fit of their optimal model. A global summary of the breakpoint distribution across time-points is also computed (e.g. to detect time-points with a large number of expression changes). Trendy does not require replicate time-points and although we focus on time-course of gene expression, Trendy may be applied to alternative features (e.g. isoform or miRNA expression) and/or other experiments with ordered conditions (e.g. spatial course).

## Implementation

Trendy is implemented in R, a free and open source language, with a vignette that provides working examples as well as an R/Shiny application to explore the fitted trends. The package and all details are publicly available at https://github.com/rhondabacher/Trendy. A visualization of the Trendy framework is given in Figure 1 and explained below.

**Figure 1.**
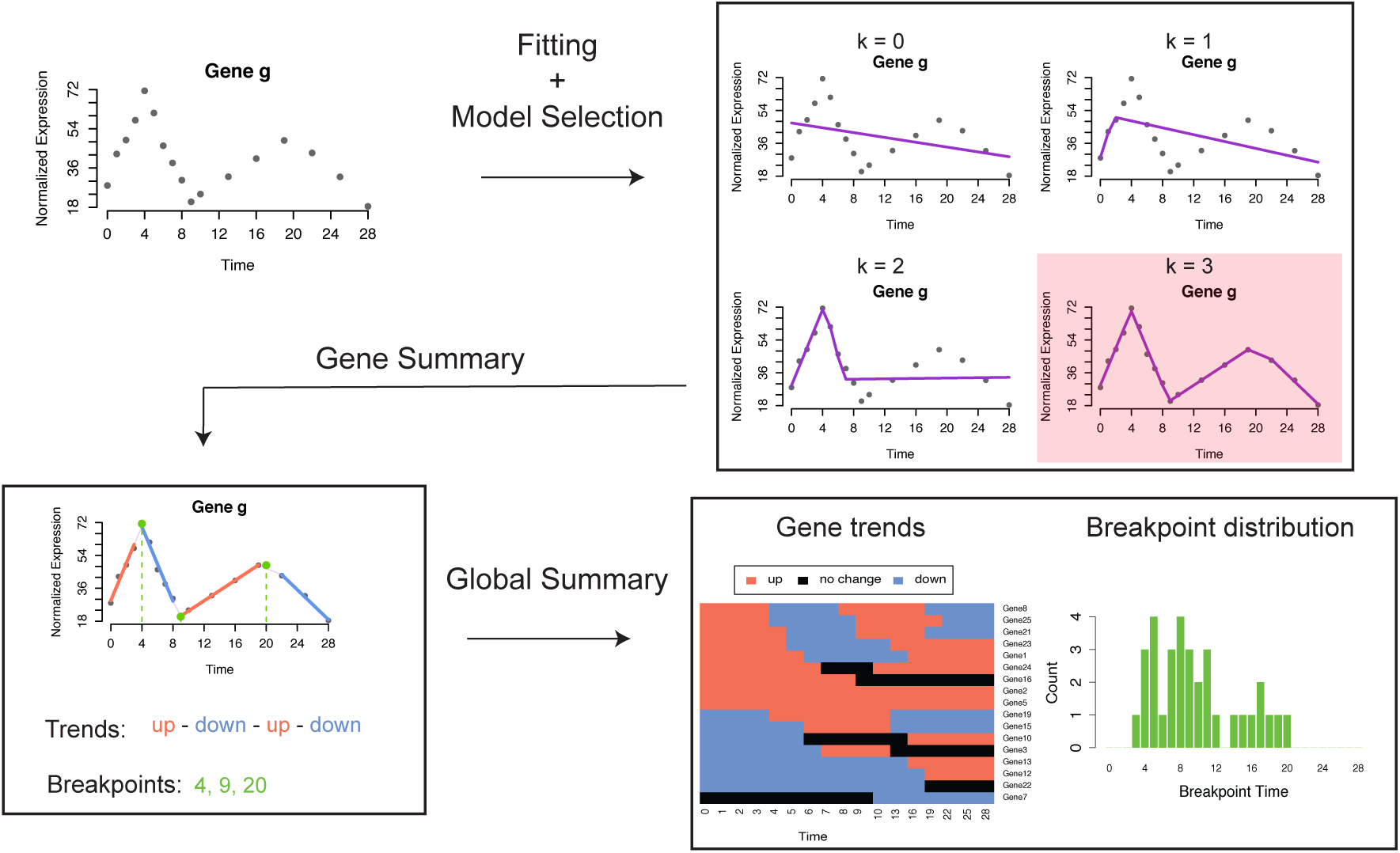
**Trendy framework**. The Trendy framework fits multiple segmented regression models to each feature/gene. The optimal model is selected as the one with the smallest BIC. Trendy summarizes the expression pattern of each gene and provides a summary of global dynamics.

### Input

The input data should be a *G* - by - *N* matrix containing the normalized expression values for each gene and each sample, where *G* is the number of genes and *N* is the number of samples. Between-sample normalization is required prior to Trendy, and should be performed according to the type of data (e.g Median-Normalization (8) for RNA-seq data or RMA (9) for microarray data). The samples should be sorted following the time-course order. A time vector, *T*, should also be supplied to denote the relative timing of each sample, this is used to specify the spacing of time-points or any replicated time-points.

### Model fitting

We denote the normalized gene expression of gene *g* and sample/time *t* as *Y*_*g,t*_. For each gene, Trendy fits a set of segmented regression models using the segmented R package (10) with the number of breakpoints varying from 1 to *K*. The default maximum number of breakpoints or changes in expression is three, but may be specified via the parameter *Max.K*.

The model for gene *g* with *k* breakpoints can be written as:

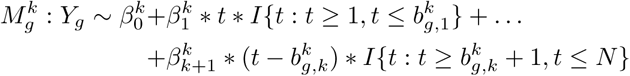

For each *k ∈* (1*, K*), the segmented regression estimates *k* breakpoints 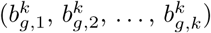 occurring between 1 and *N* and estimates *k* + 2 *β*’s. 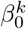 indicates the intercept and the other *β*’s indicate slopes for the *k* + 1 segments separated by the *k* breakpoints.

### Model selection

For a given gene, among the models with varying *k*, Trendy selects the optimal number of breakpoints for this gene by comparing the Bayesian information criterion (BIC) (11) for each model:

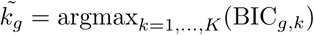

where BIC_*g,k*_ denotes the BIC for the segmented regression model with *k* breakpoints for gene *g*. To avoid overfitting, the optimal number of breakpoints will be set as 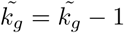if at least one segment has less than *c*_*num*_ samples. The threshold *c*_*num*_ can be specified by the user; the default is 5.

### Output

Trendy reports the following for the optimal model:

- Gene specific adjusted *R*^2^ (penalized for the chosen value of *k*): 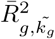
- Segment slopes: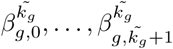
- Breakpoint estimates: 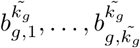

Trendy further considers only the top dynamic genes, defined as those whose optimal model has high 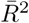, to further characterize expression patterns. The breakpoint distribution is computed along the time-course over all top genes by calculating the sum of all breakpoints at each time-point as:

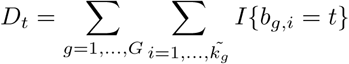

The time-points with high *D*_*t*_ can be considered as those with global expression changes.

Trendy also summarizes the fitted trend or expression pattern of top genes. For samples between the *i*^*th*^ and *i* + 1^*th*^ breakpoint for a given gene, if the t-statistic of *β*_*g,i*+1_ has p-value greater than *c*_*pval*_, the trend of this segment will be defined as no change. Otherwise the trend of this segment will be defined as up/down based on the coefficient of *β*_*g,i*+1_. The default value of *c*_*pval*_ is 0.1, but may also be specified by the user.

### Shiny application

The R/Shiny application for Trendy was developed to allow easy visualization of gene expression and the segmented regression fit. The application also allows users to extract a list of genes which follow particular expression patterns. The interface is shown in Figure 2.

**Figure 2.**
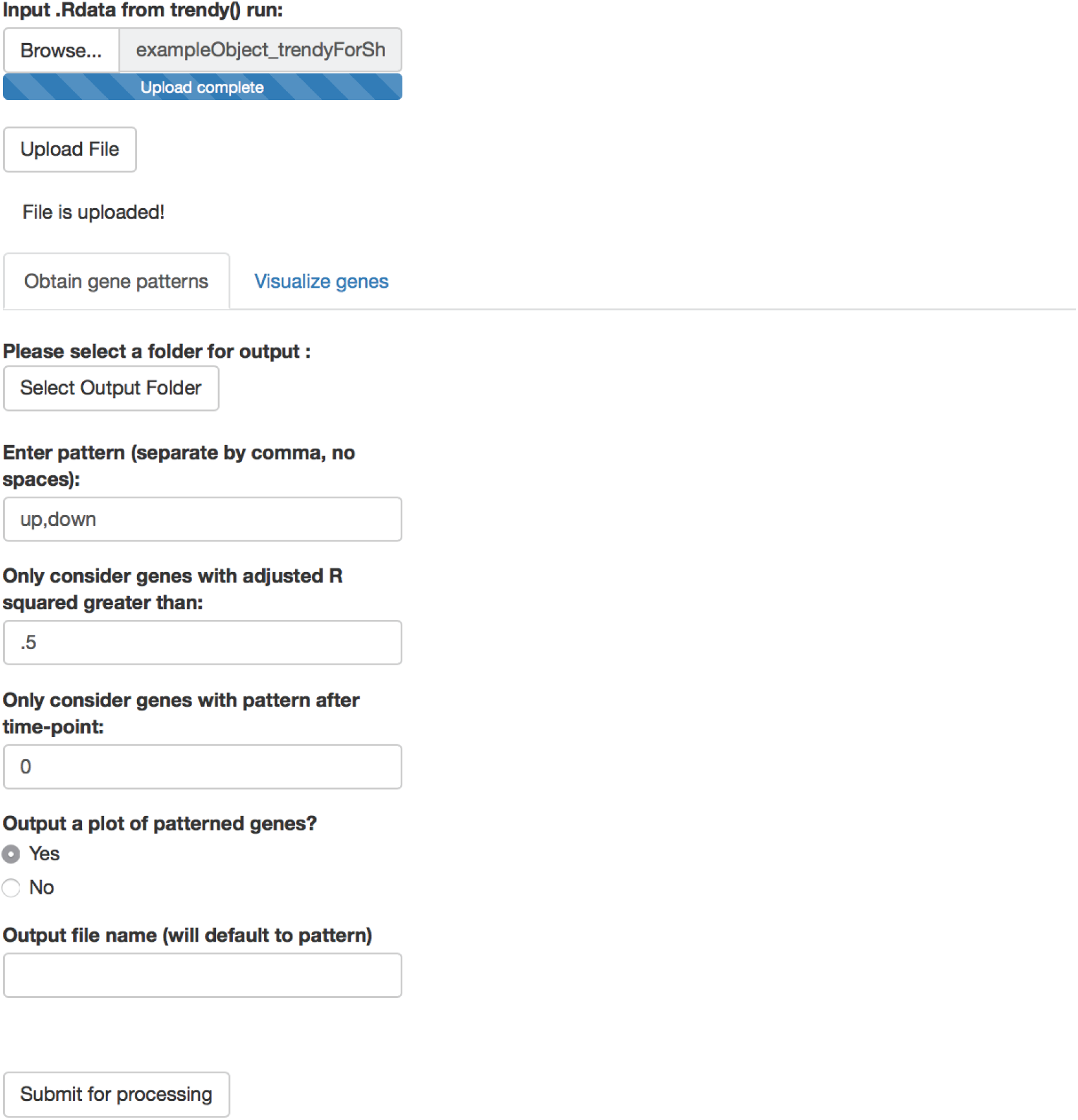
**Screenshot of Trendy R/Shiny application**. The trendy() function outputs an .RData object which can be uploaded to the Trendy R/Shiny application to explore the data and fitted trend. The user can also easily extract lists of genes with specific patterns of interest.

## Results and Discussion

We have applied Trendy to a microarray and two RNA-seq datasets datasets to demonstrate its flexibility and ability to discover relevant biological genes.

### Application to microarray data

We applied Trendy to a microarray time-course dataset from Whitfield et al., 2002(12) (experiment 3 downloaded from: http://genome-www.stanford.edu/Human-CellCycle/HeLa/). In the Whitfield data, HeLa cells were synchronized and collected periodically for a total of 48 measured time-points. Trendy identified a total of 118 top genes, defined as those having 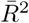*>* .8. Fig. 3(a) shows the total number of breakpoints over time for all top genes. The hours with the most breaks/changes in gene expression directly correspond to times of mitosis and completion of the cell cycle as described in Figure 1 in Whitfield et al., 2002(12). Fig. 3(b) shows two genes with fitted models from Trendy having different dynamic patterns. Both genes have 5 estimated breakpoints, however the first gene, *MAPK13*, peaks at hours 9, 22, and 34. The second gene, *CCNE1*, peaks at hours 14 and 28. These peak times also corresponds to the cell-cycle stages since CCNE1 is active during G1/S transition and MAPK13 is most active during the M phase.

**Figure 3.**
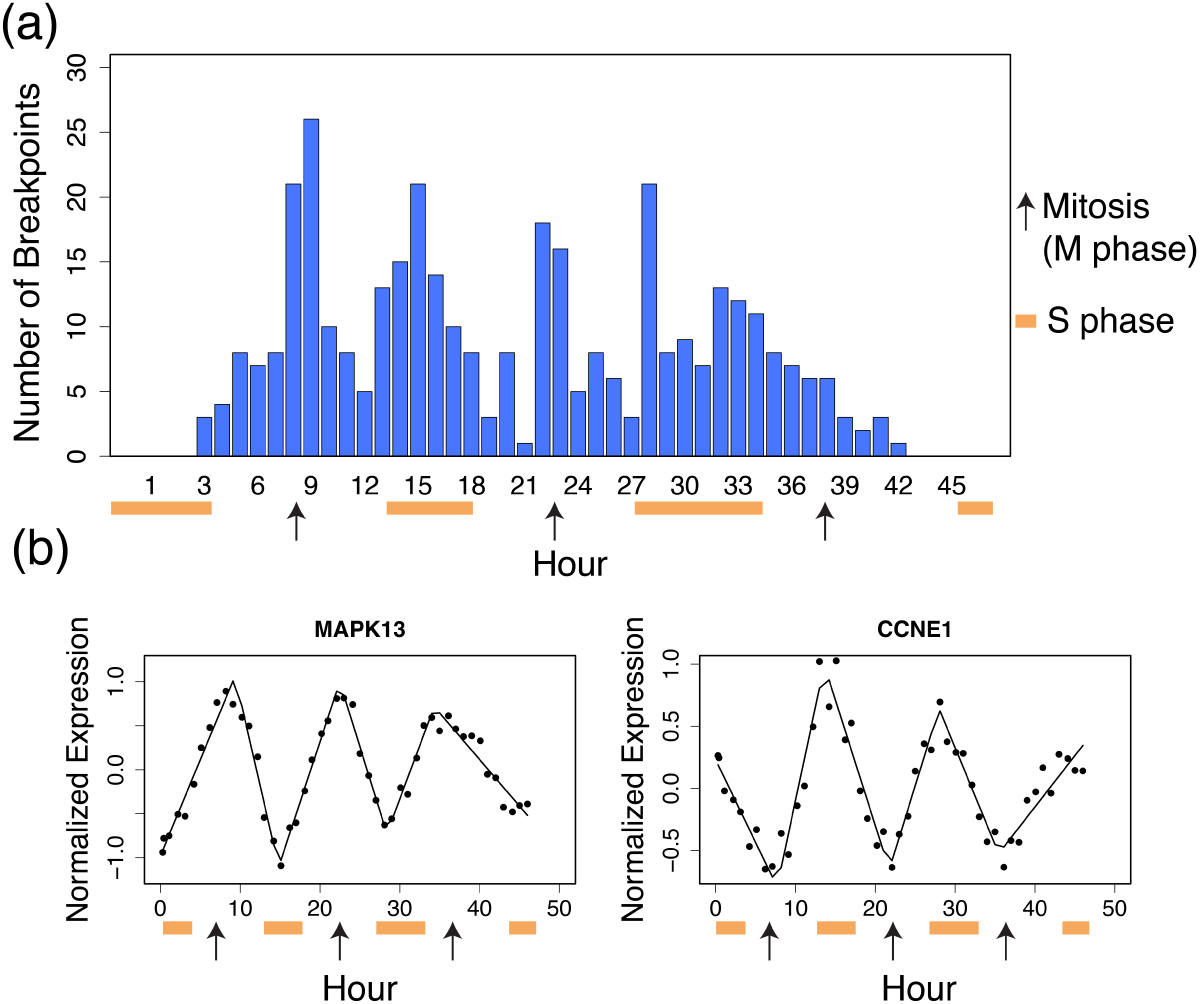
**Results of Trendy on the Whitfield dataset**. Panel (a) is the breakpoint distribution for the 118 genes having 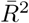*>* .8. Orange bars indicate the S phase (completion of cell cycle) and black arrows indicate the time of mitosis as shown in Figures 1 and 2 in Whitfield et al., 2002. Panel (b) contains two genes identified by Trendy with different expression dynamics over the time-course.

Further analysis by Trendy identified a total of 34 top genes that have a cycling pattern defined as “up-down-up-down” (Figure 4). Of these genes, 20 are directly annotated to the GO cell cycle pathway (GO:0007049), while others are linked to related activities such as DNA replication and chromosome organization. All but two genes were annotated to the cell cycle in the original publication, and both genes (*HBP* and *L2DTL*) are supported in the literature as being involved in the cell-cycle.

**Figure 4.**
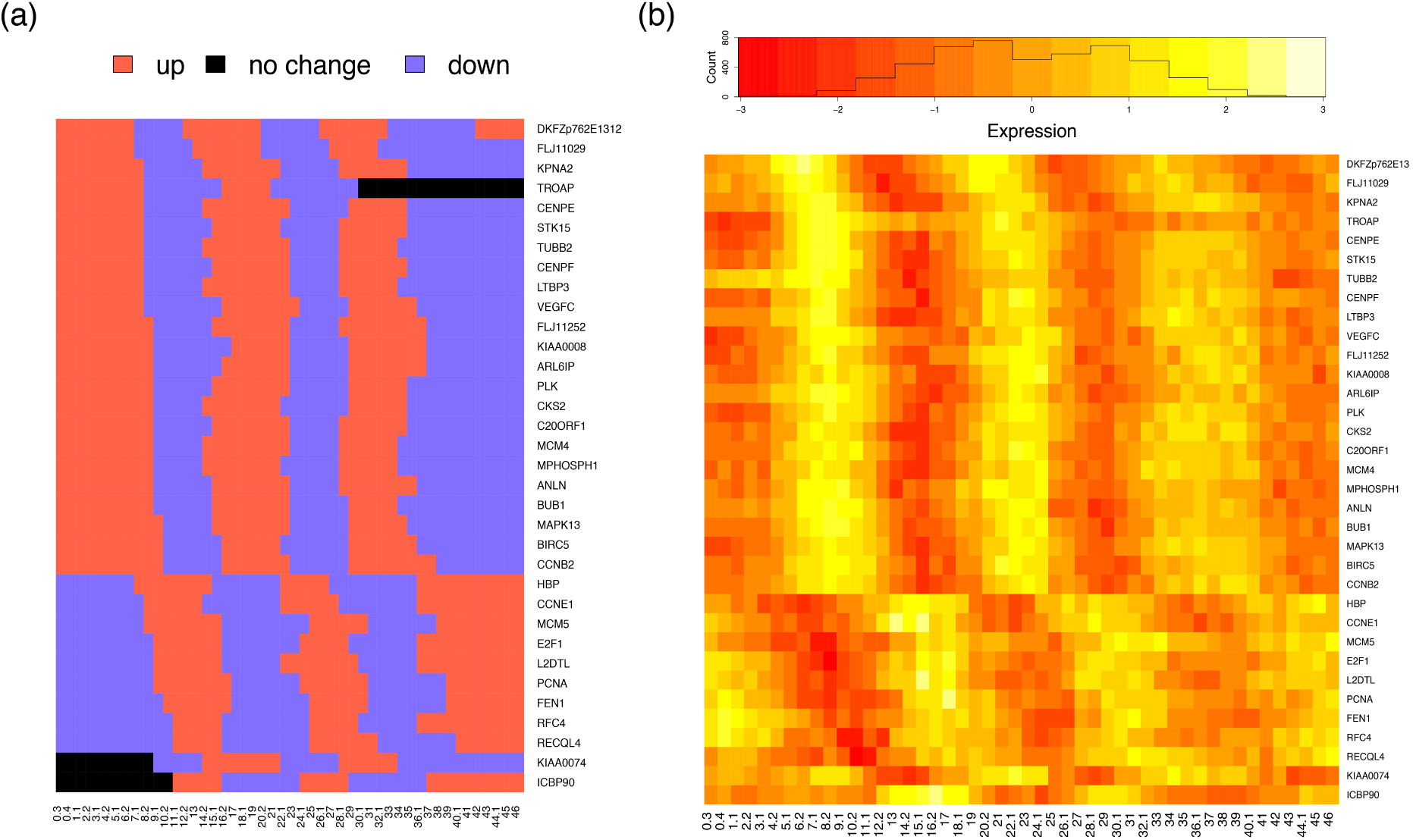
**Expression dynamics for cycling genes in Whitfield dataset**. Panel (a) shows the fitted trends for the 34 top genes having pattern “up-down-up-down” and Panel (b) show an expression heat map for the same set of genes.

### Application to RNA-seq data

We applied Trendy to the full RNA-seq time-course dataset from Jiang and Nelson et al., 2016(2) which examined axolotl embryonic development. In the axolotl data, embryos were collected at distinct developmental stages representing specific development milestones. RNA-seq was performed consecutively for Stage 1 through Stage 12, and then periodically until Stage 40 for a total of 17 stages measured. Trendy identified a total of 9,535 genes with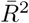 > .8. Figure 5(a) shows two genes with fitted models from Trendy having different dynamic patterns and Figure 5(b) shows the number of breakpoints over the developmental stages. The barplot appears to vary along the time-course corresponding to the waves of expression discovered in Jiang and Nelson et al., 2016(2).

**Figure 5.**
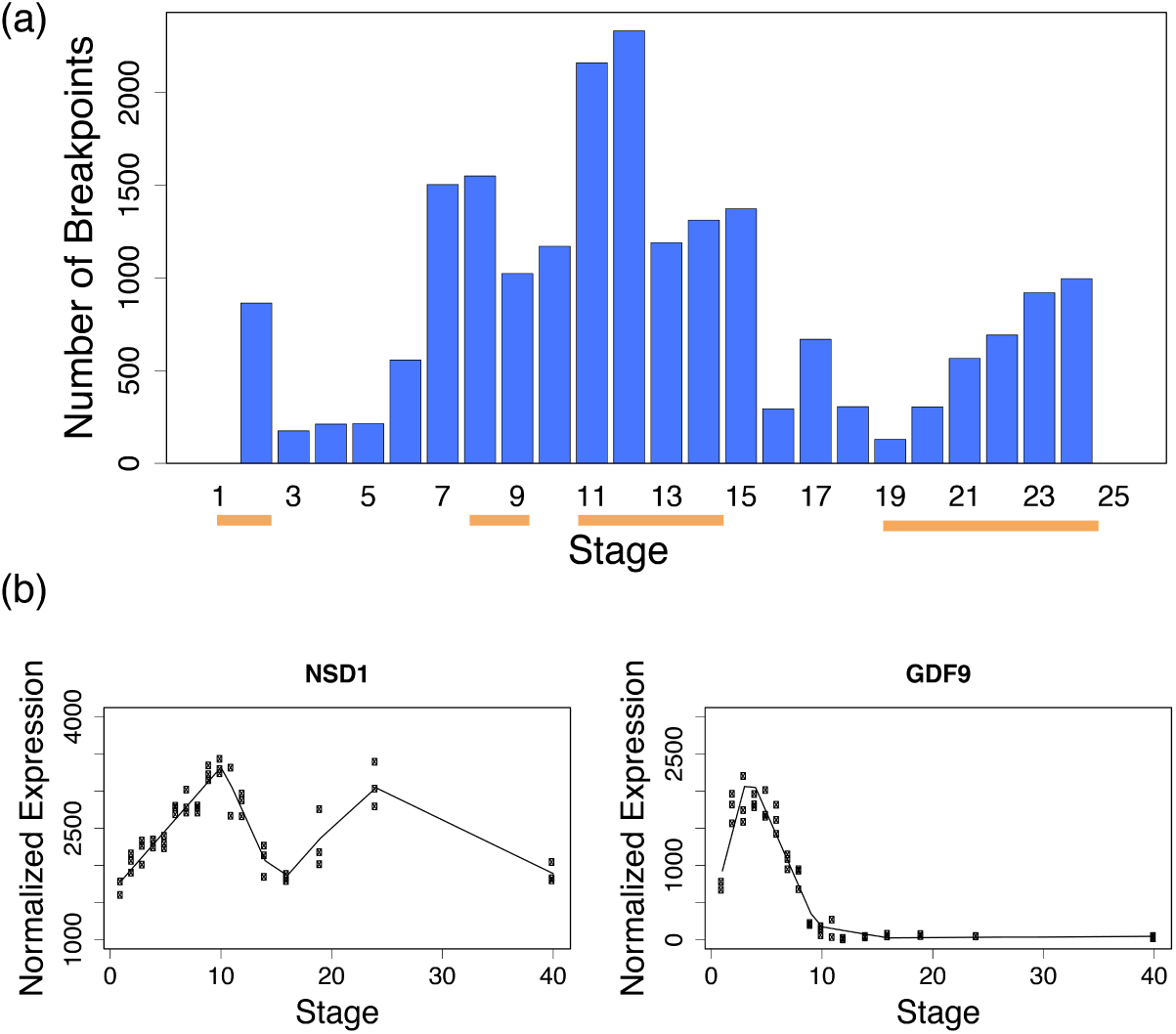
**Results of Trendy on the Axolotl dataset**. Panel (a) contains the breakpoint distribution for all 9,535 genes having 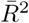 *>* .8. The orange bars indicate the times of major transcriptome changes identified in Figure 3 in Jiang and Nelson et al., 2016. Panel (b) shows two genes identified by Trendy with different expression dynamics over the time-course. The first gene, *NSD1*, has three estimated breakpoints, while *GDF9* has two breakpoints.

Further analysis by Trendy identified 807 genes having a delayed peak pattern defined as “same-up-down” with the first breakpoint occurring after Stage 8 (Figure 6). Enrichment analysis of the genes was performed based on geneset overlaps in MSigDB (v6.0 MSigDB, FDR q-value *<* .001, http://software.broadinstitute.org/gsea/msigdb)(13). The top 10 categories of enriched GO biological processes(14) include embryo development (GO:0009790), regulation of transcription (GO:0006357), organ/embryo morphogenesis (GO:0009887), tissue development (GO:0009888), regionalization (GO:0003002), and pattern specification (GO:0007389). These categories closely match those identified in the original publication(2). Genes which contain at least two peaks and appear to have cyclic activity contain enrichments for chromosome organization (GO:0051276) and regulation of gene expression (GO:0010629) within the top ten categories. The full set of enrichment results are given in Supplementary File 1.

**Figure 6.**
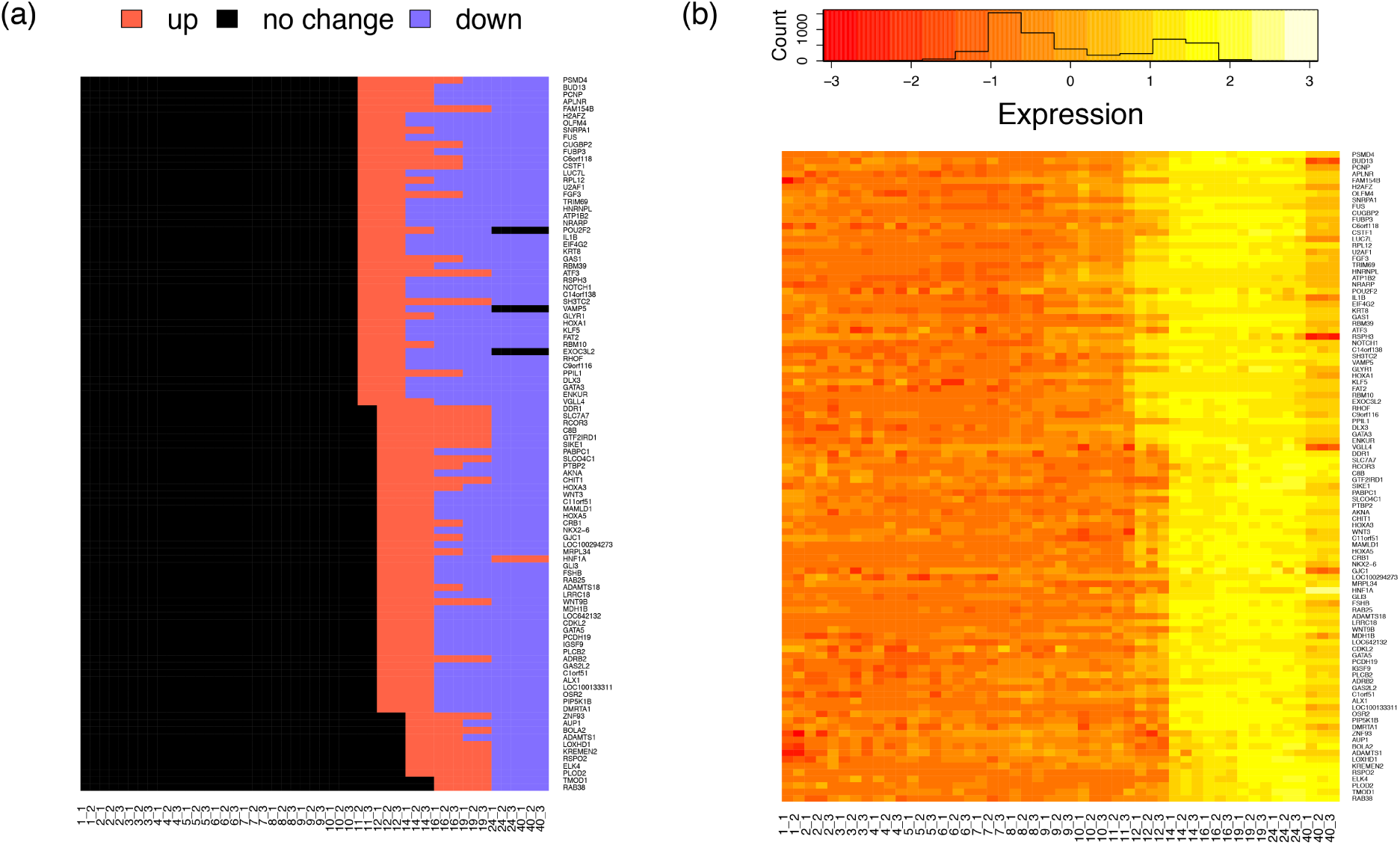
**Expression dynamics for delayed peak genes in the Axolotl dataset**. Fitted trends and expression heat maps are shown for all a subset of 100 top genes having pattern “same-up-down” wth breakpoint after Stage 8.

Trendy was also applied to two neural differentiation time-course RNA-seq experiments in Barry et al., 2017(1). Breakpoints were estimated separately for the mouse and human differentiation time-course experiments and peaking genes were identified as those having the pattern “up-down”. The authors found that the relationship between mouse and human peak-times for top ranked neural genes closely matched that expected by the gold-standard Carnegie stage progressions (1).

### Conclusion

The prevalence of experiments containing ordered conditions is on the rise with increases in bulk time-course sequencing experiments to study dynamic biological processes, in addition to the proliferation of single-cell snapshot sequencing experiments in which cells can be computationally ordered and assigned a temporal (or spatial) order (Leng et al., 2015b; Chu et al., 2016; Trapnell et al., 2014).

We developed an R package, Trendy, to analyze expression dynamics in high throughput profiling experiments with ordered conditions. Trendy provides statistical analyses and summaries of feature-specific and global expression dynamics. In addition to the standard workflow in Trendy, also included in the R package is an R/Shiny application to visualize and explore expression dynamics for specific genes and the ability to extract genes containing user-defined patterns of expression.

## Acknowledgments

The authors would like to thank Chris Barry for helpful feedback regarding the formatting of output from Trendy.

### Author’s contributions

RB and NL created the Trendy package under the guidance of JT, CK, and RS. RB conducted the data analysis. LFC contributed guidance and interpretation of results. RB and NL wrote the manuscript. All authors read and approved the final manuscript.

